# MYC directly transactivates CR2/CD21, the receptor of the Epstein-Barr virus, and enhances the viral infection of Burkitt lymphoma cells

**DOI:** 10.1101/2022.12.28.522143

**Authors:** Ester Molina, Lucia Garcia-Gutierrez, Vanessa Junco, Mercedes Perez-Olivares, Virginia G. de Yébenes, Rosa Blanco, Laura Quevedo, Juan C Acosta, Ana V. Marín, Daniela Ulgiati, Ramon Merino, Ignacio Varela, José R. Regueiro, Ignacio Moreno de Alborán, Almudena Ramiro, Javier León

## Abstract

The molecular hallmark of Burkitt lymphoma (BL) is a chromosomal translocation that results in deregulated expression of *MYC* oncogene. This translocation is present in virtually all BL. MYC is an oncogenic transcription factor deregulated in about half of total human tumors, by translocation or other mechanisms. Transcriptomic studies reveal more than 1000 genes regulated by MYC but a much smaller fraction of these genes is directly activated by MYC. All the endemic BL and many sporadic BL cells are associated to the Epstein-Barr virus (EBV) infection. The currently accepted mechanism for the MYC and BL association is that EBV is the causing agent inducing MYC translocation. Complement receptor 2 or CR2 (also called CD21) is a membrane protein that serves as EBV receptor in lymphoid cells. Here we show that CR2 is a direct MYC target gene. This conclusion is based on several evidences. First, MYC downregulation is linked to *CR2* downregulation both in proliferating and in arrested cells. Second, MYC binds human *CR2* promoter and this binding depends on E-box elements. Third, MYC activates *CR2* promoter in an E-box dependent manner. Four, MYC activates *CR2* transcription in the absence of protein synthesis. Importantly, MYC also induces *CR2* expression in mouse primary B cells. Thus, CR2 is a *bona fide* MYC direct target gene. Moreover, higher MYC expression levels in Burkitt lymphoma-derived cells result in a more efficient EBV infection. We propose an alternative mechanism compatible with the correlation between EBV infection and MYC translocation observed in endemic BL, i.e., that deregulated MYC in BL cells occurs first and favors the EBV infection.

## INTRODUCTION

MYC is a transcription factor that belongs to the b-HLH-LZ family of proteins and binds DNA as a dimer with MAX, another b-HLH-LZ protein. MYC-MAX heterodimers bind to the DNA through their basic domain to consensus sequences called E-boxes (CANNTG consensus motif), present in most MYC target genes [1, 2]. MYC transforms cells modulating functions essential in tumor cell biology as cell cycle progression energetic metabolism, lipid and nucleotide biosynthesis, protein synthesis, apoptosis and differentiation block, among others [1-3]. In line with its wide array of functions, MYC regulates more than one thousand genes [4-7]. However, the identification of direct MYC target genes is hampered because MYC is a weak transactivator but provokes potent changes in cell physiology. Therefore, many of these genes are regulated as a secondary effect of proliferation, anaerobic metabolism, apoptosis, etc. Indeed, only a minority of genes (3-48% depending on the particular study, mean 21% from 9 independent reports) with MYC bound at the promoter are identified as direct MYC target genes (reviewed by [8]).

MYC is one of the most commonly activated oncogenes in human cancer, being deregulated in about 50% of human malignancies of different types [9, 10] and it is particularly prevalent in lymphoma and leukemia [11]. MYC deregulation occurs through different mechanisms, including translocations, gene amplifications and other less understood mechanisms that result in increased *MYC* transcription [2, 12]. High grade B cell lymphomas show a high prevalence of MYC deregulation, usually associated to worse prognosis [13]. Among them, Burkitt lymphoma (BL) is an aggressive lymphoma that was originally described in Africa equatorial [14]. It is the most common childhood cancer in areas were malaria is endemic in Africa but also in Brazil and Papua New Guinea. This BL variant is called endemic BL (eBL) and affects mainly children with high prevalence (5-10 per 10^5^ /year). Clinically and histologically similar BL occurs in Europe and USA affecting adults with much lesser prevalence, a variant called sporadic BL (sBL). A third less common variant is the immunodeficiency-associated BL [15, 16].

BL is believed to originate in the germinal center of lymph nodes. In the germinal center, rapidly dividing B cells undergo somatic hypermutation and class-switch recombination of the immunoglobulin heavy chain genes. MYC plays a prominent role in germinal center formation [17]. Activation-induced cytidine deaminase (AICDA) initiates somatic hypermutation and class switch recombination [18], but it can also induce Ig/MYC translocations [19]. *MYC* is translocated to one of the immunoglobulin *loci* in nearly all BL cases, resulting in aberrant *MYC* expression under the control of the immunoglobulin enhancers. The most frequent translocation occurs into the immunoglobulin heavy chain [t(8;14)(q24;q32)] but it could also be found in the light chain *loci* [t(2;8)(p11;q24) and t(8;22)(q24;q11)] [13, 20, 21] Indeed, transgenic mice carrying an Ig enhancer linked to MYC (Eμ-MYC mice) develop aggressive B cell lymphomas with high penetrance [22]. Besides translocations, 30-50% of BL also carry mutations within the MYC gene [23], most of them generating a more stable MYC protein [24].

Epstein-Barr virus (EBV) is a herpesvirus that was originally discovered associated to endemic BL where is present in more than 90% of the cases. In contrast, EBV is detected in 1-2% of adults and 30-40% of children with sporadic BL and 30-40% of patients with the immunodeficiency-associated BL [15, 25]. EBV establishes a harmless lifelong infection in B cells in over 95% of adults worldwide, and it is associated to infectious mononucleosis [26] and multiple sclerosis [27, 28]. EBV was the first virus described with oncogenic potential due to its association to BL and its ability to transform B cells into immortalized lymphoblastoid cell lines [29]. EBV can display three gene expression programs, called latency programs I, II and III respectively, in which different sets of viral genes are expressed [25, 27, 29]. These genes are involved in processes leading to B cell transformation, such as immortalization [30], resistance to apoptosis [31] and metabolic reprogramming [32].

The most accepted hypothesis to explain the high incidence of EBV in endemic BL proposes a causative role for EBV. This hypothesis stemmed from the finding of high EBV antibodies titers in sera of 12 out of 16 African children before the BL diagnosis [33]. This was reproduced in subsequent reports and leading to the declaration of EBV as a causative agent of BL by the World Health Organization [34]. EBV, malaria and HIV infection, are cofactors that may cooperate with MYC in promoting B cell proliferation in the germinal center by reducing apoptosis or facilitating the immune evasion of the tumor [15, 16]. However, the pathogenic mechanism by which EBV genes promote MYC translocation and cooperate with MYC in BL tumorigenesis is unclear [27, 35]. Furthermore, some data do not fit with the hypothesis of EBV being the cause of BL. EBV infection is absent in a majority of the sporadic BL cases, while EBV infection is associated to other tumors such as nasopharyngeal carcinoma, gastric cancer and other hematological malignancies (Hodgkin leukemia, post-transplant lymphoproliferative disease, T cell lymphoma) where MYC translocation is absent (reviewed in [25, 27, 35]). Strikingly, BL cells express the latency type I viral genetic program which includes only a protein, EBNA1, plus two small non coding nuclear RNAs, EBER1, EBER2, and several miRNA. However, *EBNA1* is involved in EBV episomal replication but is not oncogenic. Actually, BL cells are the only EBV-infected cells showing this limited viral genome expression [27, 29]. EBV infects B lymphocytes through a membrane glycoprotein, the complement receptor 2 or CR2 (also termed CD21) receptor [36]. CR2 acts as a co-receptor for B Cell Receptor (BCR) and is found in a complex with other proteins such as CD19 or CD81 [37].

In this work, we show that MYC directly induces CR2 expression in BL-derived cell lines and primary B cells, which contributes to the increased proliferation mediated by MYC. Induction of CR2 by MYC leads to higher number of EBV-infected cells overexpressing MYC. Based on this, we propose an alternative (but not mutually exclusive) mechanism in which the upregulation of MYC due to chromosomal translocation increases the virus receptor density within the B cell surface, leading to B-lymphocytes prone to infection by EBV.

## RESULTS

### MYC downregulation leads to decreased CR2 expression in lymphoma cells

Previous studies on gene expression profiling of K562 leukemia cell line with conditional MYC expression revealed CR2 as one of the genes regulated by MYC (J.C. Acosta, PhD Thesis Dissertation, University of Cantabria, 2005). Given the role of CR2 as EBV receptor and the role of MYC in BL, we investigated whether CR2 could be a novel MYC target gene. First, we treated Raji cells (BL cell line) and Jurkat cells (acute T cell leukemia cell line) with JQ1, an inhibitor of the bromodomain and extra terminal (BET) family of proteins which is known to potently inhibit MYC transcription in leukemia and lymphoma cells [38]. JQ1 treatment downregulated MYC expression at the mRNA level in Raji and Jurkat cells, and this was accompanied with a decrease in CR2 mRNA expression (Figure 1A). The decrease in MYC and CR2 expression was also detected at the protein level in both cell lines (Figure 1B). MYC is necessary to maintain cell cycle progression and MYC downregulation is known to arrest proliferation [3]. In agreement, treatment of Raji and Jurkat cells with increasing concentrations of JQ1 decreased their proliferative capacity in a dose-dependent manner as shown by cell counting (Supplementary Figure 1A) and increased the percentage of cells in G^1^ phase (Supplementary Figure 1B). Longer exposure to JQ1 resulted in apoptosis, as shown by annexin V staining in Raji and Jurkat cells (Supplementary Figure 1C).

**Figure 1.**
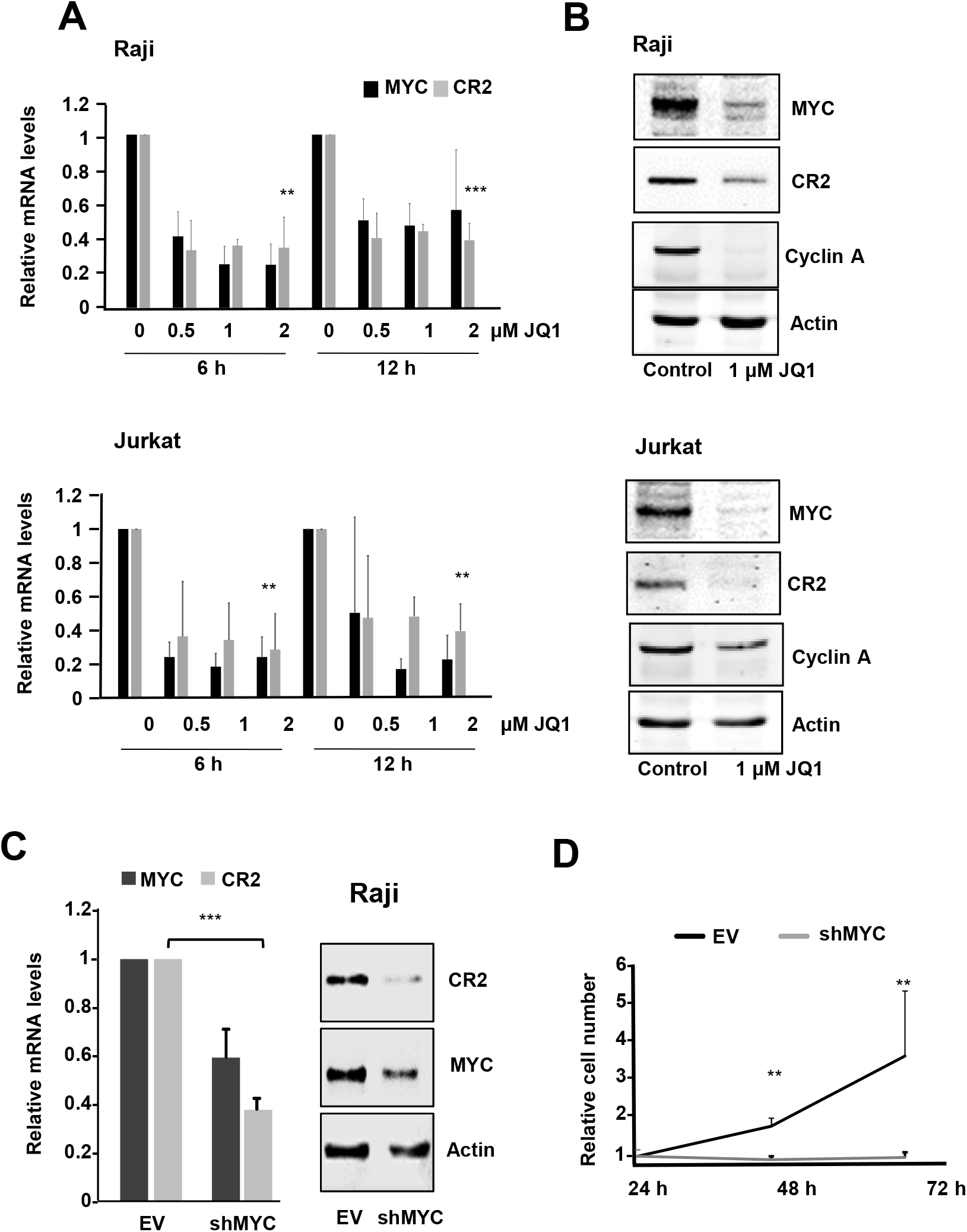
Down-regulation of MYC results in lower CR2 expression. (**A**) mRNA expression of MYC and CR2 in Raji and Jurkat cells treated with JQ1 at the indicated concentrations for 6 and 12 h. The expression was determined by RT-qPCR. Data represent mean values ±S.D. (n = 3). ** *P* < 0.005; *** *P* < 0.001. (**B**) Immunoblot showing the protein levels of MYC and CR2 in Raji and Jurkat cells treated with JQ1 at the indicated concentrations for 24 h. β-Actin levels were used loading control. **(C)** MYC and CR2 mRNA (left panel) and protein (right panel) levels assayed by RT-qPCR and immunoblot, respectively, from Raji cells infected with shMYC-containing lentiviral particles and selected with puromycin. β-actin levels were used as loading control. (**D**) Proliferation of Raji cells transduced with shMYC lentiviral particles and control virus. Data represent mean values ±SD (n=3), normalized to cell counting at the start of the experiment. \*\**P* <0.05; ***P < 0.005

To investigate a direct effect of MYC downregulation on *CR2* expression, we asked whether *MYC* silencing with short-hairpin (sh) RNA constructs leads to *CR2* downregulation. We knocked down MYC expression in Raji cells using shMYC-containing lentiviral particles (a mixture of two sh constructs targeting human MYC). The results showed that MYC depletion significantly reduced the levels of CR2 mRNA and protein were significantly reduced upon lentiviral infection compared to controls (Figure 1C). The rate of cell proliferation was also dramatically reduced upon MYC silencing (Figure 1D).

### MYC upregulates *CR2* mRNA expression independently of cell proliferation

The former results can be explained if CR2 expression is linked to the proliferation state of the cell, so it would decrease as a consequence of MYC-mediated proliferation arrest, rather than by direct transcriptional regulation by MYC. To address this issue, we used KMycJ cells, a K562 derived cell line with ectopic MYC expression inducible by Zn^2+^ addition [39]. In these cells, JQ1 treatment decreases endogenous *MYC* expression while the addition of Zn^2+^ induced the exogenous MYC (Figure 2A). However, the arrest in proliferation of KMycJ cells upon JQ1 treatment was not restored by ectopic MYC expression (Figure 2B). Importantly, the induction of MYC by addition of Zn^2+^ resulted in the upregulation of *CR2* mRNA upregulation even in arrested cells (Figure 2C). Total MYC levels did not increase in growing cells treated with Zn^2+^ likely due to a well-known MYC auto suppression mechanism by which exogenous MYC downregulates endogenous *MYC* levels in most systems including hematopoietic cell lines, so that total MYC levels do not raise above a certain threshold [40, 41]. Despite the original claim that JQ1 (and derivatives) inhibits proliferation primarily through MYC downregulation, these cells were arrested by JQ1 even in the presence of high MYC levels. However, an increase in *CR2* expression was observed upon MYC induction by Zn^2+^ in JQ1 arrested cells (Figure 2B,C). To confirm that CR2 regulation by MYC is independent from the proliferative state of the cells, we performed a similar approach by inducing cell cycle arrest in KMycJ cells using 12-O-tetradecanoylphorbol-13-acetate (TPA), a drug known to arrest proliferation and dramatically reduce MYC levels in K562 cells [41]. Cells were treated with TPA and then the ectopic MYC expression was induced with Zn^2+^. As shown in Figure 2D, TPA induced a proliferative arrest that was not rescued by MYC upon Zn^2+^ addition. Importantly, *CR2* mRNA levels increased upon MYC induction even in conditions of arrested proliferation (Figure 2E).

**Figure 2.**
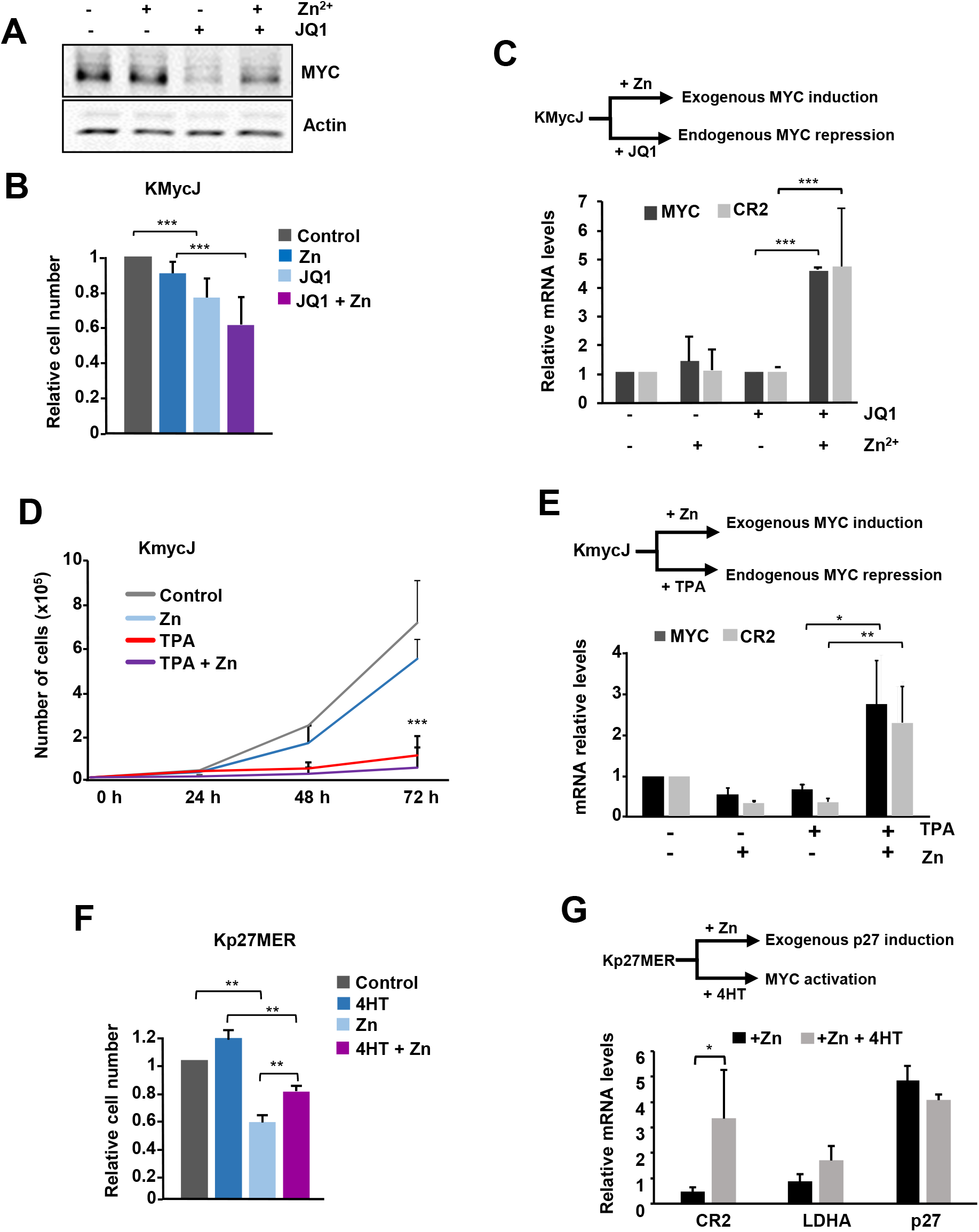
MYC induces CR2 expression in arrested cells. **(A)** Immunoblot showing MYC protein levels of Raji cells treated for 24 h with 75 μM ZnSO_4_ and 1 μM JQ1 determined by immunoblot. β-actin levels were used as loading control. **(B)** Proliferation of KMycJ cells treated with 75 μM ZnSO_4_ and 1 μM JQ1 for 48 h, measured by cell counting and normalized to untreated cells. Data represent mean values ±SD (n = 4). **(C)** Up: scheme of the conditional MYC expression by Zn^2+^ and repression by JQ1 in KmycJ cells. Lower graph: *MYC* and *CR2* mRNA levels in KMycJ cells treated for 24 h with 75 μM ZnSO_4_ and 1 μM JQ1, quantified by RT-qPCR. Data represent mean values ±SD (n=2). **(D)** Proliferation of KMycJ cells pretreated with TPA and 12h later treated with Zn^2+,^ measured by cell counting. Data represent mean values ±SD (n = 3). **(E)** Up: scheme of the conditional MYC expression by 75 μM ZnSO_4_ and repression by 10 nM TPA in KmycJ cells. Lower graph: *MYC* and *CR2* expression levels in KMycJ cells treated with TPA and/or Zn^2+^ for 48 h determined by RT-qPCR. Data represent mean values ±SD (n=2). **(F)** Proliferation of Kp27MER cells treated with 200 nM 4HT and/or 75 μM ZnSO_4_ for 48 h, measured by cell counting and normalized to untreated cells. Data represent mean values ±SD (n = 4). ** P < 0.01. **(G)** Up: scheme of the conditional MYC activation by 4HT and induction by Zn^2+^. Lower graph: expression of *CR2, p27/CDKN1B* and *LDHA* expression from Kp27MER determined by RT-qPCR. Data represent mean values ±S.D. (n=4); * P < 0.05.

As a parallel approach to investigate whether MYC induced CR2 in non-growing cells, we used another K562 derived cell line, Kp27MER, which contains a *CDKN1B* transgene (p27^KIP^) inducible by Zn^2+^ and the chimeric protein MYC-ER which becomes activated by 4-hydroxy-tamoxifen (4HT). p27^KIP^ is a CDK and cell cycle inhibitor and as expected, addition of ZnSO_4_ induced p27 expression and a concomitant decrease in cell proliferation after 48 h, which is rescued only partially by MYC-ER activation (Figure 2F). However, despite this proliferation arrest, we observed an increase in *CR2* expression after MYC-ER activation (Figure 2G). As a control we confirmed the MYC-dependent upregulation of *LDHA*, a *bona fide* MYC target gene [42]. Taken together, the results suggest that *CR2* induction is a direct effect of MYC and not a consequence of the proliferative state of the cells.

### MYC directly binds to CR2 promoter

Chromatin immunoprecipitation (ChIP)-seq data generated by the ENCODE Consortium (UCSC database; https://genome-euro.ucsc.edu, GRCh37/hg19) predicted two peaks of MYC and MAX occupancy within the proximal region of CR2 promoter in human B cells (Figure 3A). Bioinformatic analysis revealed that there are four canonical E-boxes in the region surrounding the CR2 transcription start site (TSS), two of which mapped within the promoter near the TSS and the other two within the first intron (Figure 3B). To confirm MYC binding to CR2 promoter in BL derived cells we performed ChIP with an anti-MYC antibody in Raji cells using different primers along the CR2 proximal promoter and within the first intron. The results showed an enrichment of MYC binding to the proximal region of CR2 promoter in a region containing two E-box clustered within a 20 bp sequence located upstream the TSS of *CR2* (Figure 3C). This is consistent with our hypothesis that MYC is a regulator of *CR2* transcription.

**Figure 3.**
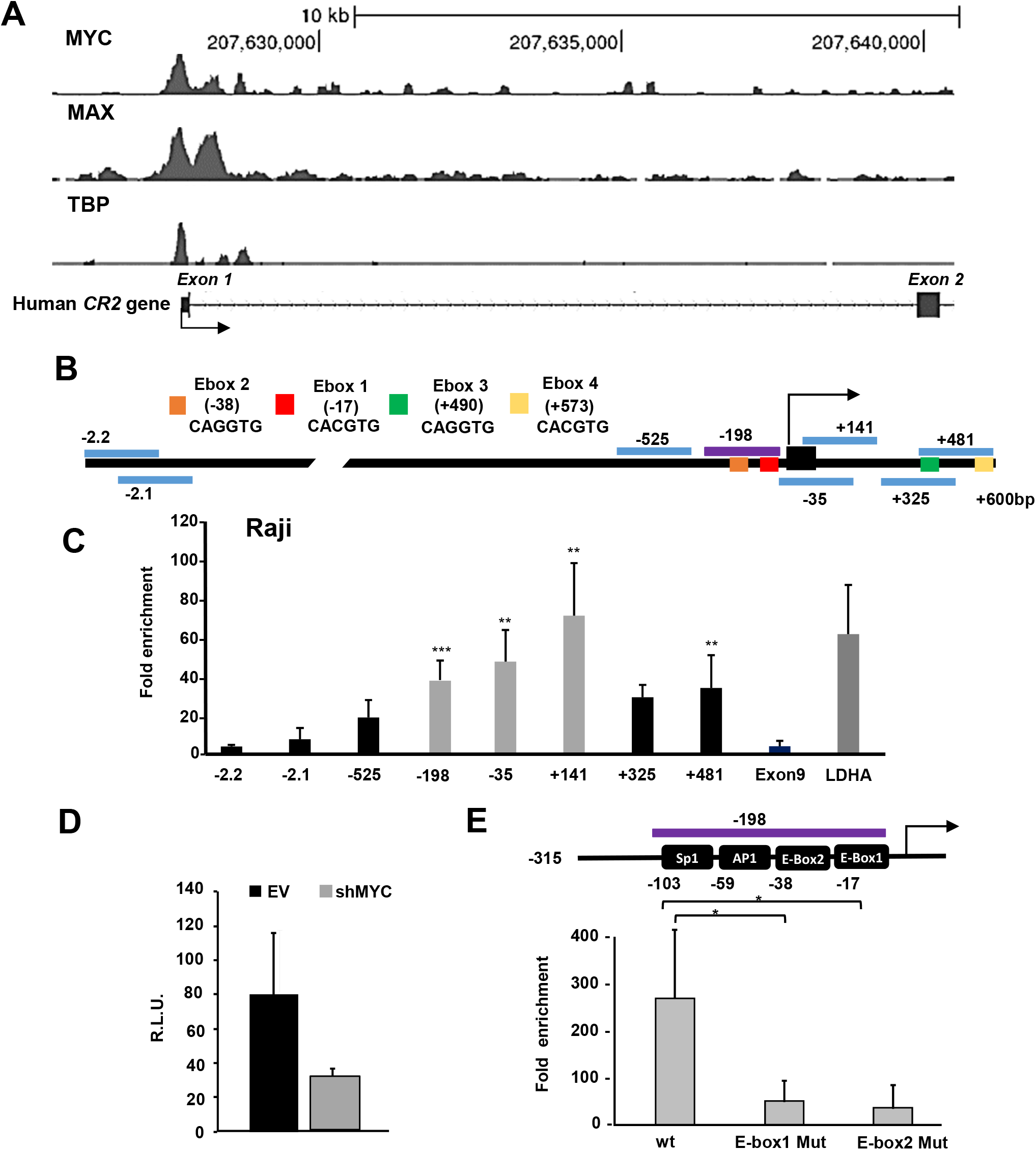
Binding of MYC to the promoter of CR2. (**A**) Schematic representation of *CR2* chromosome localization and MYC and MAX binding sites according with ChIP-seq data of the ENCODE project (USC Genome Browser, GRCh37/hg19hg19 release) in EBV-transformed human B cells (GM12878 cell line). TBP (TATA-box binding protein) is also shown to mark the transcription initiation complex binding. (B) Schematic representation of CR2 proximal promoter and the first exon and intron. The horizontal bars represent the region amplified by the different primers used in the ChIP analysis. (**C**) ChIP with anti-MYC antibody on CR2 gene of Raji cells. Exon9 of CR2 gene was used as a negative control and a LDHA proximal promoter region mapping -85 bp to +19 bp was used as positive control for MYC binding. Data are mean values, relative to IgG, ±SD (n=3). (**D**) CR2 promoter activity measured by luciferase assay of Raji cells transfected with a *CR2* promoter luciferase reporter and transduced with shMYC-containing lentiviral particles (or the corresponding empty vector, EV). Luciferase activity was determined 48 h after infection. Data represent mean values ±SD (n=3). (**E**) MYC occupancy on *CR2* promoter assayed by ChIP using an anti-MYC antibody in 293T cells transfected with the luciferase constructs containing either wild type or mutated E-box1 or E-box2 *CR2* promoter regions as indicated. wt, wild type promoter. The amplicon analysed is shown schematically above. Data show relative MYC fold enrichment in each sample with respect to IgG. Data represent mean values ±S.D. (n = 3) * P < 0.1; ** P < 0.05; *** P < 0.01.

To validate the MYC-dependent transactivation of *CR2* we performed luciferase assays with the luciferase reporter gene under the control of the CR2 promoter region harboring the two E-boxes located ∼200 bp upstream the TSS. The reporter was transfected along with a construct for short-hairpin MYC. The results showed that knocking down MYC expression lead to a dramatic drop of the *CR2* promoter activity (Figure 3D). To assess the relevance of the E-boxes located within the CR2 promoter for MYC transactivation we carried out ChIP assays to study MYC occupancy on two reporter constructs containing deletions in either E-box 1 or E-box 2. The results showed that the lack of any of the two E-boxes significantly decreased MYC occupancy as compared with that of the wild type CR2 promoter (Figure 3E). These results show that MYC binds CR2 promoter and stimulates its transcription in an E-box-dependent manner. Therefore, we conclude that *CR2* is *bona fide* MYC target gene.

### MYC-ER activation upregulates CR2 expression in the absence of *de novo* protein synthesis

Our results showed that MYC induces CR2 expression independently on MYC-mediated effects on proliferation. However, these results do not rule out the possibility that another MYC-target gene encoding for a different transcription factor could be mediating CR2 up-regulation. To test whether CR2 is a direct MYC target, we took advantage of a K562 derived cell line engineered with the MYC-ER chimeric protein (KMycER cells) activatable by 4HT [39]. To determine whether MYC was capable of inducing *CR2* expression in the absence of *de novo* protein synthesis we treated KMycER cells with cycloheximide, a protein synthesis inhibitor. This model would allow discriminating between a direct MYC transcripcional effect on CR2 promoter form an indirect mechanism (Figure 4A). Endogenous MYC protein levels dramatically decreased after 6 h of cycloheximide treatment, as expected, while activation of MYC-ER by the addition of 4HT stabilized the chimeric protein (Figure 4B). The treatment with 4HT, which activated MYC-ER, increased *CR2* mRNA levels even in cycloheximide-treated cells, where new proteins cannot be synthesized, ruling out the possibility of an intermediate MYC-target gene regulating CR2 expression (Figure 4C). Taken together these data show that MYC directly activates CR2 transcription.

**Figure 4.**
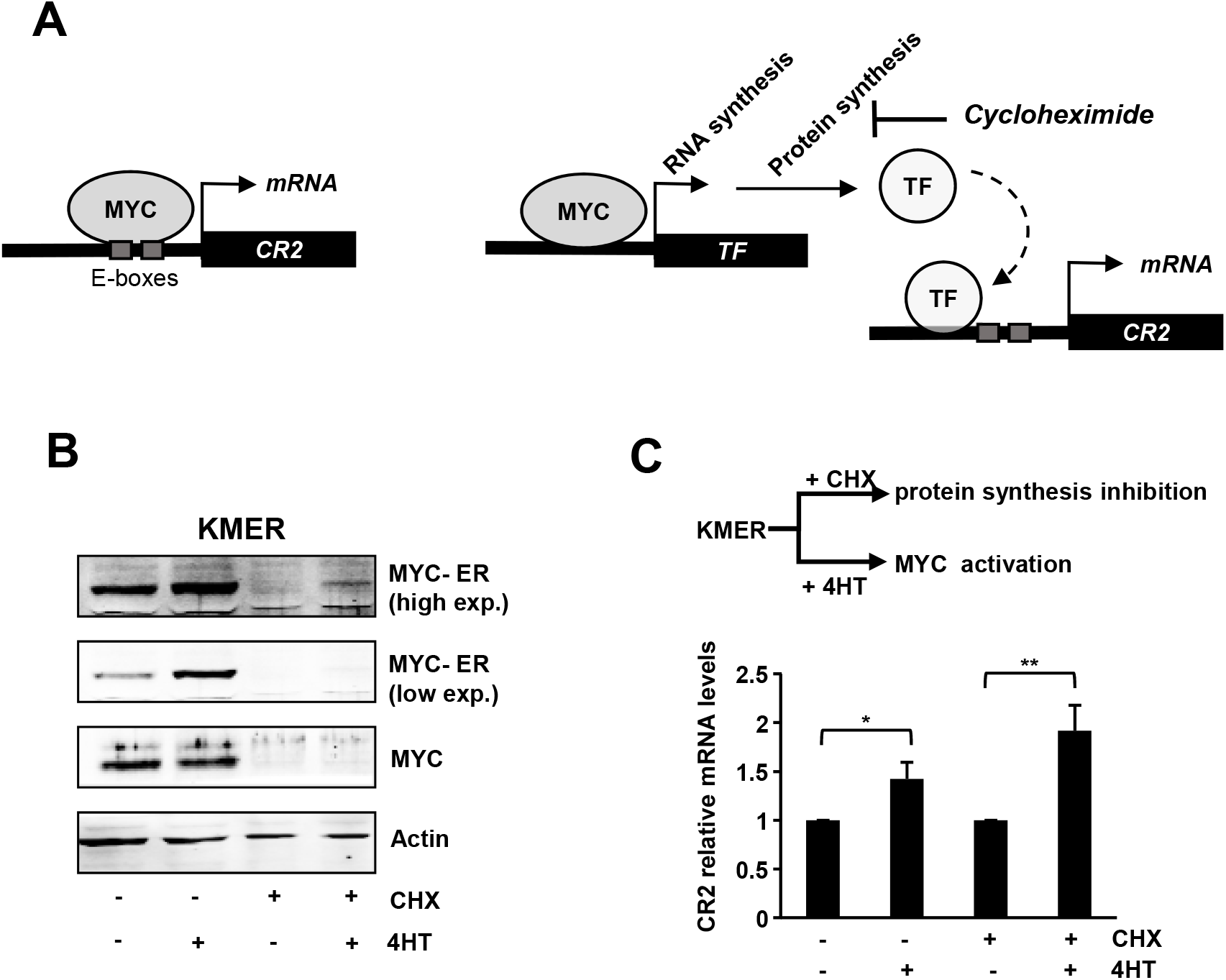
CR2 is a direct MYC target gene. **(A)** Scheme showing the two possible mechanisms from MYC-mediated induction of *CR2* expression. Left: the direct cycloheximide insensitive mechanism. Right: the indirect cycloheximide sensitive mechanism by which MYC would induce another transcription factor (TF) which in turn would activate *CR2* expression. **(B)** MYC and MYC-ER protein levels from KMER cells treated with 200 nM 4HT and 50 μg/ml cycloheximide (CHX). After 6 h of treatment, cell lysates were prepared and analyzed by immunoblot. β-actin levels were used as loading control. **(C)** Upper part: scheme of the conditional MYC activation by 4HT and effect of cycloheximide (CHX) in KMER cells. Lower graph: expression of *CR2* mRNA in KMER cells treated with 4HT and CHX. Data represent mean values ±S.D. (n = 3). * P < 0.05; ** P < 0.005.

Interestingly, CR2 interacts with the B cell receptor (BCR) complex [43]. and contributes to the proliferative signal transduction from the receptor. Indeed, it has been reported that CR2 is required for optimal B cell proliferation [44, 45]. Thus, we asked whether MYC-mediated upregulation of CR2 can also contribute to B cell proliferation in our model system. To explore this, we silenced *CR2* through lentiviral constructs expressing short hairpin constructs in Raji cells (Supplementary Figure 2A) and assayed the effect of CR2 depletion on proliferation. The results showed that CR2 knockdown produced a dramatic decrease in the proliferation of Raji cells (Supplementary Figure 2B).

### MYC induces CR2 expression in primary B cells

To address whether MYC induces CR2 expression also in primary B cells, we analyzed by flow cytometry the expression of MYC and CR2 in splenic B lymphocytes from C57/BL6 wild type mice. We found that the fraction of cells with 10% higher MYC expression levels also showed higher CR2 expression (Figure 5A), thus confirming the correlation observed in human cell lines. A correlation between MYC and CR2 mRNA levels can also be found in hematopoietic human tissues (Gene Expression Profiling Interactive Analysis, http://gepia.cancer-pku.cn/) (Supplemental Figure 3). To confirm this correlation we analyzed CR2 expression in mice where *Myc* can be conditionally depleted in primary B lymphocytes in the *Myc*^*fl*/fl^;*Max*^*fl*/+^;*Cd19*^*cre*/+^;*Rosa26*^*gfp*/gfp^ (MycKO-Cd19) model previously described by us [46]. This mouse carries one allele of *Max* and both alleles of *Myc* flanked by *loxp* sites. Expression of Cre recombinase is driven by the endogenous promoter of *Cd19* and promotes specific *Myc* deletion in B lymphocytes and expression of GFP [46]. GFP allows the rapid identification and analysis of B lymphocytes that have undergone Cre-mediated deletion of *Myc*. Primary B lymphocytes from the spleens of MycKO-Cd19 and heterozygous control mice were activated with LPS and interleukin-4, stained with anti-CR2/CD21 antibody and analyzed by flow cytometry. We observed a dramatic decrease in the population of CR2/CD21^+^ cells within the GFP^+^ population (*Myc* deleted cells) in the MycKO-Cd19 compared to heterozygous control mice before and after 48 h of activation (Figure 5B). To see whether this effect was specific for CD21, we analyzed the surface expression of the activation marker CD69 on the same cells by flow cytometry. We did not see any significant differences of CD69 surface expression between mutant and control cells (Supplementary Figure 4).

**Figure 5.**
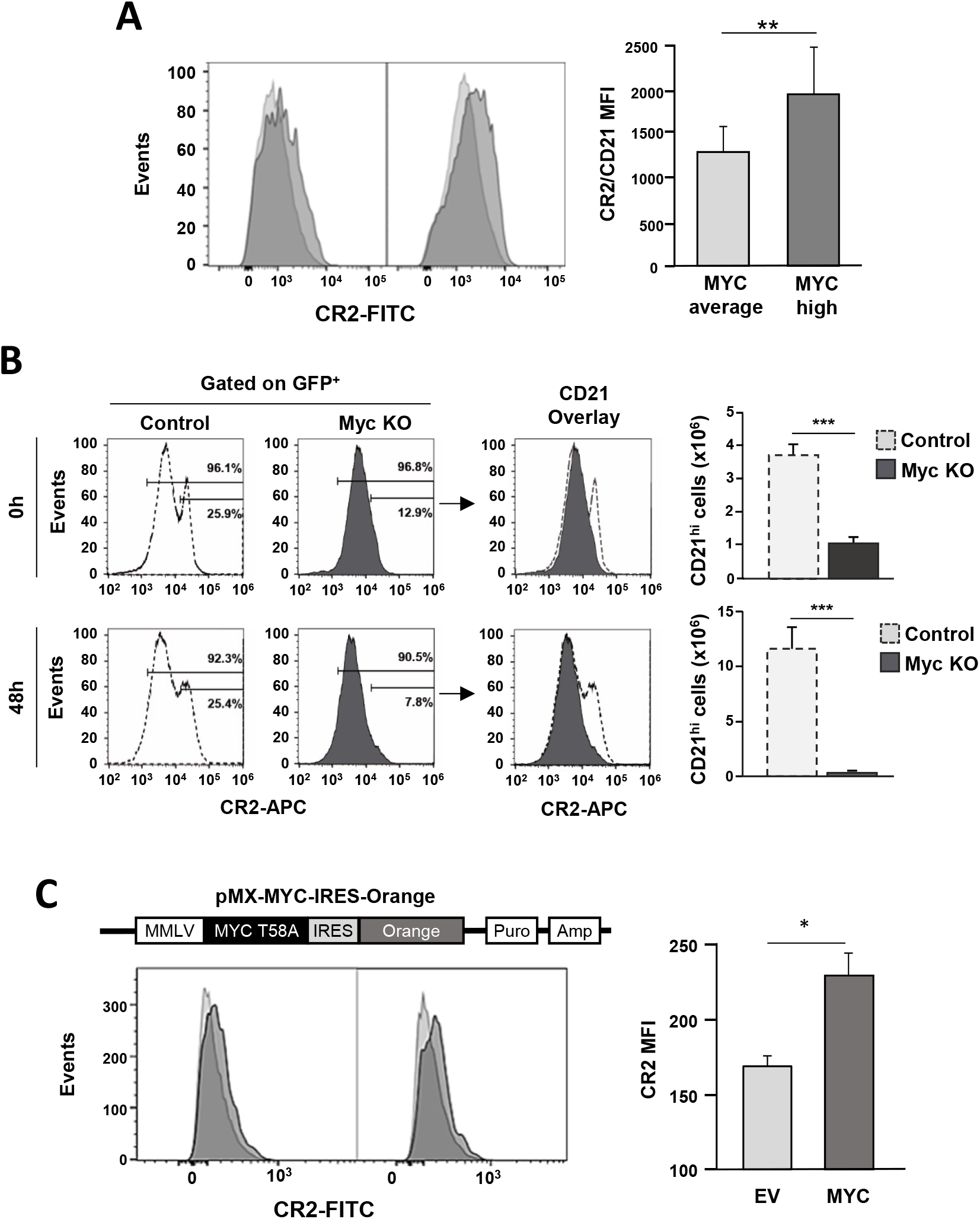
Myc induces CR2 expression in primary B cells. **(A)** Cell surface CR2/CD21 levels in splenic mouse B cells with the 10% highest MYC expression levels (dark grey) as compared to cells the rest of the cells (light grey). B cells were prepared from 18 weeks old C57/BL6 wild type mice analyzed by flow cytometry. Two representative overlay histograms are shown at the left and quantification of CD21 mean fluorescence intensity (MFI) at the right. Data are mean values ± S.D. (n = 6) **p < 0.005. **(B)** Mature B lymphocytes from the spleens of *Myc*^*flox/flox*^*;Max*^*flox/+*^*;cd19*^*cre/+*^*;Rosa26*^*gfp/gfp*^ (MycKO, *n*=3) homozygous and *Myc*^*flox/+*^*;Max*^*flox/+*^*;Cd19*^*cre/+*^*;Rosa26*^*gfp/gfp*^ heterozygous control (*n* = 3) mice were activated with LPS and IL-4 for 48h. Cells were stained with the indicated antibodies and analysed by flow cytometry. GFP^+^ cells (*Myc* deleted B lymphocytes) were gated and surface expression of CR2/CD21 was analysed. Absolute numbers of CD21^hi^ are shown. *n* = 3, \*\*\**p*<0.001. **(C)** Left panel, up: schematic representation of the pMX-MYC-IRES-Orange retroviral vector used in the experiments. MMLV: LTR and ψ sequences of MMLV virus; Puro and Amp, puromycin and ampicillin resistance genes. Right panel, bottom: expression of CR2 in mouse splenic B cells after MYC overexpression. CD43^-^ B cells isolated from spleens were cultured with LPS and interleukin-4 for 24h and transduced with pMX-MYC-Orange retrovirus and the control retrovirus (EV) during 48 h. CR21 was assayed in Orange+ transduced cells. Two representative overlay histograms are shown at the left and quantification of CD21 mean fluorescence intensity (MFI) at the right Data are mean values ±S.D., n = 5, \**p* < 0.05.

To confirm the induction of CR2 by MYC, we overexpressed MYC in primary mouse splenic B cells by retroviral transduction with MYC, encoded in an Orange-reporter bicistronic IRES vector (Figure 5C), as well as with control Orange retrovirus without MYC. We found that MYC-transduced cells expressed higher CR2 levels (Figure 5C). In light of these data we conclude that MYC promotes *CR2* expression in murine primary B cells.

### MYC depletion decreases EBV infection efficiency

Our results indicate that the EBV receptor, CR2, is a direct MYC target gene. Thus, we wanted to elucidate whether the modulation of MYC expression correlated with the infection capacity of EBV. Unlike Raji cells, Ramos is an EBV-negative BL cell line and thus a suitable model to test this hypothesis. We confirmed first that gDNA from Raji cells contained EBV genes while gDNA from Ramos cells did not, as expected (Figure 6A). We next induced partial MYC depletion in Ramos through shMYC lentiviral transduction and confirmed that *MYC* (left panel) and *CR2* (right panel) expression levels were downregulated at mRNA level (Figure 6B) compared with cells infected with empty vector. To test the correlation between MYC expression and EBV infection, we decreased *MYC* levels by shMYC and infected Ramos cells with EBV-containing supernatants from a producer B95-8 cell line. After 48 h, total genomic DNA (gDNA) was prepared from control and MYC-depleted Ramos cells and two EBV genes, LMP1 and EBNA1, were measured to quantify the levels of EBV infection by using PCR. We observed that cells previously infected with shMYC-containing lentivirus showed reduced content of EBV genes within their genome (Figure 6C, D). These results cannot be explained by a delayed growth of MYC-silenced cells as the proliferation rate of the cells did not significantly modify the extent of EBV infection (Supplementary Figure S5).

**Figure 6.**
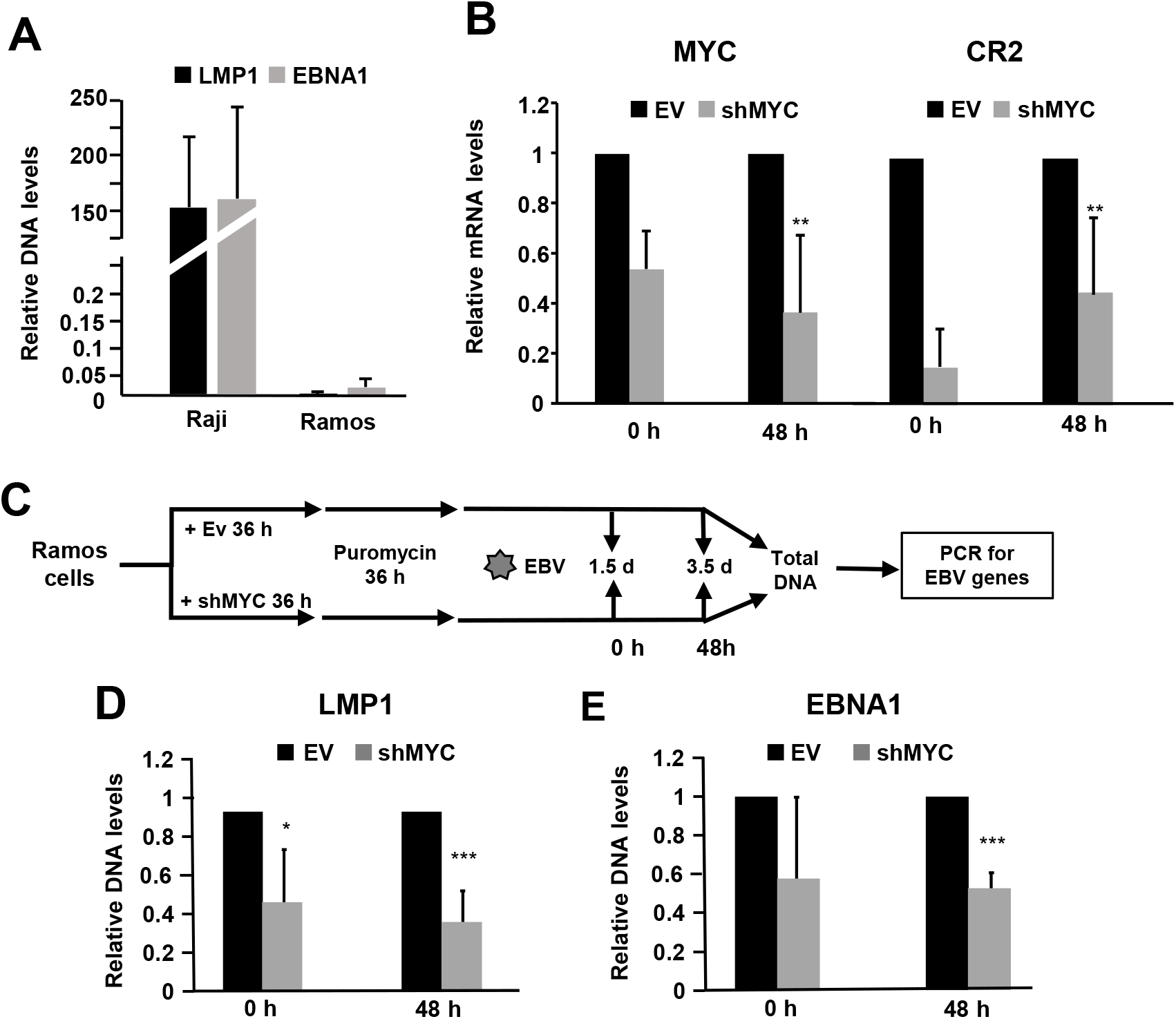
MYC depletion decreases EBV infection of B cells. (**A**) DNA levels of LMP1 and EBNA1 viral genes from Raji and Ramos cells infected with EBV (supernatant from B95-8 cells) line analyzed by PCR. Data normalized against *LDHA* and *CR2* DNA levels (same genomic regions used in ChIP experiments (Supplementary Table S1). (**B**) *MYC* (left graph) and *CR2* (right graph) mRNA levels from Ramos cells infected with lentiviral particles expressing shMYC and empty vector (EV) and analyzed by RT-qPCR. (**C**) Scheme of the experiment. (**D, E**). *LMP1* and *EBNA1* genomic DNA levels in Ramos cells infected with EBV, quantified by qPCR. Data represent mean values ±S.D. (n = 3) ** *P* < 0.05; *** *P* < 0.005.

In a second approach we infected Ramos cells with a lentiviral vector that constitutively expressed MYC (LvMYC), as shown in the Supplementary Figure S6C). We infected Ramos cells with this lentvirus. Three days after lentiviral infection and puromycin selection, cells were infected with EBV (B95-8 cells supernatants) for further 48 h (Figure 7A). The protein levels of MYC in infected cells was confirmed by immunoblot (Figure 7B). Total gDNA was prepared and the levels of EBNA1 and LMP1 DNA were analyzed by PCR. The results showed an increased infection in cells expressing higher MYC levels (Figure 7C). Altogether, the results indicate that EBV infection’s ability is dependent, at least partially, on MYC levels.

**Figure 7.**
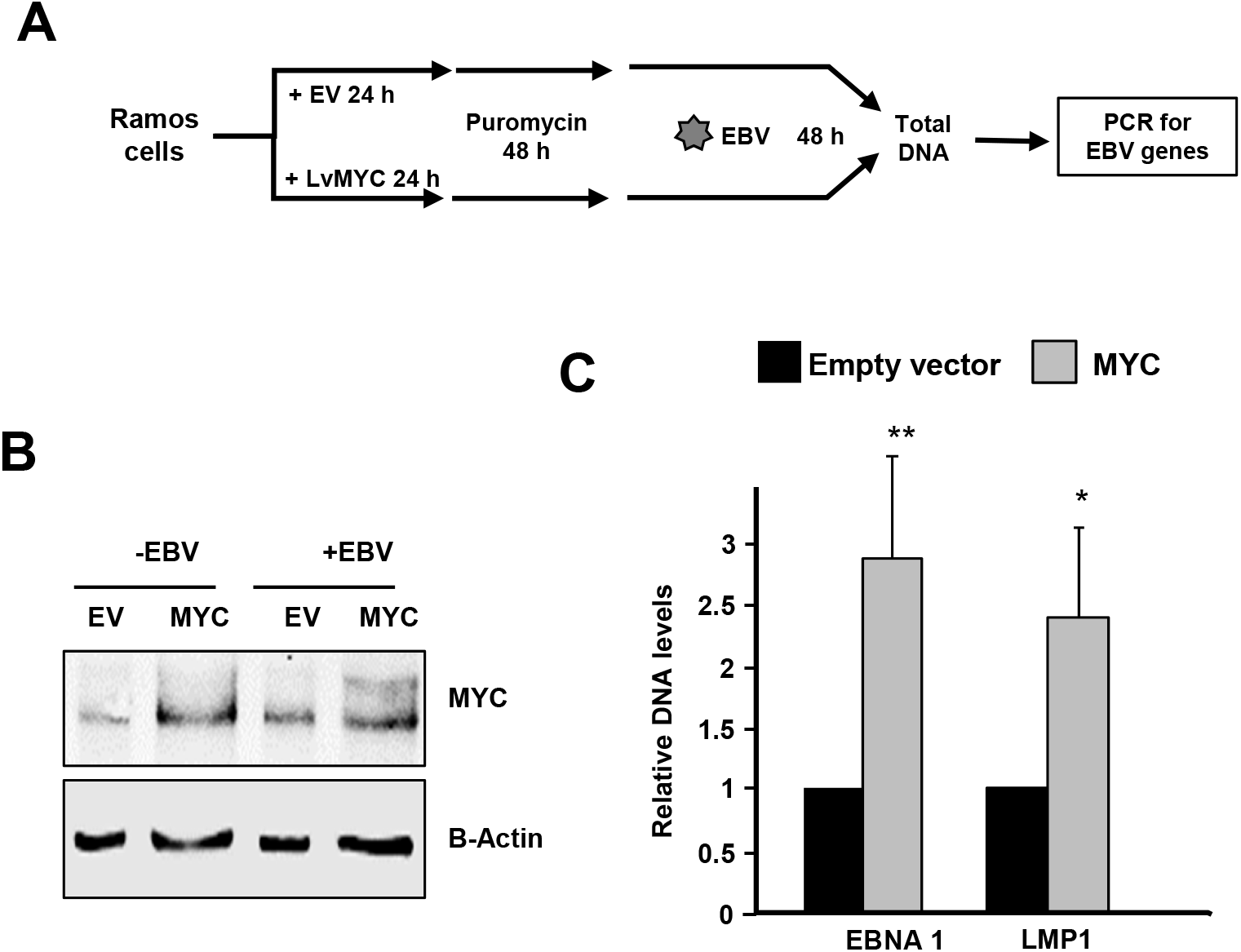
MYC overexpression increases EBV infection of B cells. (**A**) Scheme of the experiment. (**B**) Immunoblot showing the levels of MYC in cells infected with lentivirus expressing MYC and the empty vector (EV) after 48 h of EBV infection. Actin was used as loading control. (**C)**. LMP1 and EBNA1 genomic DNA levels in Ramos cells infected MYC lentivirus and EBV, quantified by qPCR. Data represent mean values ±S.D. * *P* < 0.05 (n = 3).

## DISCUSSION

In this work we propose a new mechanism for the pathogenesis of Burkitt lymphoma, based on the MYC-mediated of the CR2 Epstein-Barr virus receptor. In our initial transcriptomic studies in human leukemia cells, we observed an increased expression of CR2 upon *MYC* enforced expression. We confirmed this observation in B and T cell lines (Raji and Jurkat cell lines respectively) with the drug JQ1, described as a potent inhibitor of MYC expression in B cells, and with short hairpin (sh) constructs against MYC. We found that MYC downregulation induced by either JQ1 or shMYC is accompanied by a decrease in CR2 expression. These results, together with our initial observation, confirmed that MYC regulates CR2 expression, a result also observed in transcriptomic studies in different cell lines including B cells [47] [6, 48]. As expected from the low MYC levels, both JQ1 and shMYC caused proliferation arrest. Therefore, it was conceivable that CR2 downregulation could be a consequence of the proliferation arrest instead of being directly related to the transactivation activity of MYC. The same effect may apply to other MYC regulated genes revealed in transcriptomic studies. To analyze MYC induction of CR2 in the absence of cell proliferation we used human leukemia cells engineered with inducible MYC alleles and observed *CR2* up-regulation in response to MYC expression even in arrested cells upon JQ1 treatment. Noteworthy, JQ1 and other BET inhibitors are currently being tested in clinical assays in B cell malignancies primarily due to their effect on MYC expression [38]. Our results show that ectopic MYC expression fails to rescue the anti-proliferative effect of JQ1. Thus, our study provides additional evidence that the anti-tumoral effect of the BRD4 inhibitor is not only dependent on MYC repression. Additionally, two independent models in which MYC activity can be induced upon arrested proliferation i.e. by TPA treatment or p27 enforced expression, showed that MYC-dependent CR2 induction is not a consequence of the MYC-induced proliferation.

These results showed that *CR2* upregulation after MYC overexpression (or *CR2* downregulation after MYC depletion) is independent from the proliferative state of the cells, consistently with *CR2* being a direct MYC target gene. To confirm this we performed (i) ChIP experiments showing MYC binding close to the human *CR2* transcription start site, which required the presence of two proximal E-boxes and (ii) luciferase reporter assays showing that MYC activated the *CR2* promoter in a E-box dependent manner. These results strongly suggested that CR2 is a direct target gene, but it must be noted that first, binding to chromatin does not mean that the gene is actually transactivated by a transcription factor, and second, that there are many transcription factors that are MYC target genes. Thus, it is possible that MYC induces other transcription factor(s) which in turn activate(s) the CR2 promoter. We addressed this point with cell lines expressing the MYC-ER chimera (activatable by 4HT) and using the protein synthesis inhibitor cycloheximide. With this system we showed that MYC induced the expression of CR2 mRNA in the absence of *de novo* protein synthesis. Thus, we conclude that *CR2* is a direct MYC target gene in B cells. However, the analysis of the expression data from 60 BL available in public databases (cBioPortal) shows that there is not a tight correlation between MYC and CR2 levels, although MYC missense mutations are significantly more frequent in lymphomas with higher CR2 expression (http://cbioportal.org) [49]. It is possible that CR2 upregulation by MYC occurs at the first stages of cell transformation into BL cell and is not maintained in subsequent advanced stages. Also, there are other transcription factors known to regulate *CR2* expression [50-52].

The relationship between EBV infection and MYC translocation events in BL is a matter of debate. The most accepted hypothesis proposes that EBV infection would lead to the activation and expansion of germinal center B cells and subsequent MYC translocation (schematized in Figure 8). Indeed, the correlation between high EBV antibody titers and BL found in children living in Africa led to declare EBV as the causative agent of BL [34]. However, the mechanisms whereby EBV leads to MYC deregulation remains ill-defined and other evidence argues against this “virus first” model. First, in most sporadic BL cases EBV infection is absent while *MYC* is translocated in all BL cases. Moreover, although there are differences in the prevalence of some alterations, there are not differential molecular markers in the mutational landscape and transcriptome between EBV^+^ and EBV^-^ lymphomas [53-55]. Second, more than 95% of the human adult population has been infected by EBV, but oncogenic events related with EBV infection have a very low incidence. Third, *MYC* translocation is not found in other tumors associated with EBV infection, remarkably nasopharyngeal cancer, gastric cancer and Hodgkin lymphoma [56-58]. Fourth, the EBV genes required for B cell transformation in lymphoblastoid cell lines are not expressed in BL. Indeed, the latency I genetic program expressed in BL (i.e., the EBNA1 protein and EBER1-2 non-coding RNAs) is unable to transform B cells. It has been reported that EBNA2 [59, 60] and EBNA3C [61] induce MYC expression, and that EBNA3C induces activation-induced cytidine deaminase activity, which could generate chromosomal translocations involving MYC [19, 62, 63] and likely marks other genetic changes in EBV-infected BL [55]. However, neither EBNA2 nor EBNA3C are expressed in BL. Moreover, MYC represses the EBV transforming LMP1 gene [64] and inhibits the reactivation of the viral lytic cycle [65].

**Figure 8.**
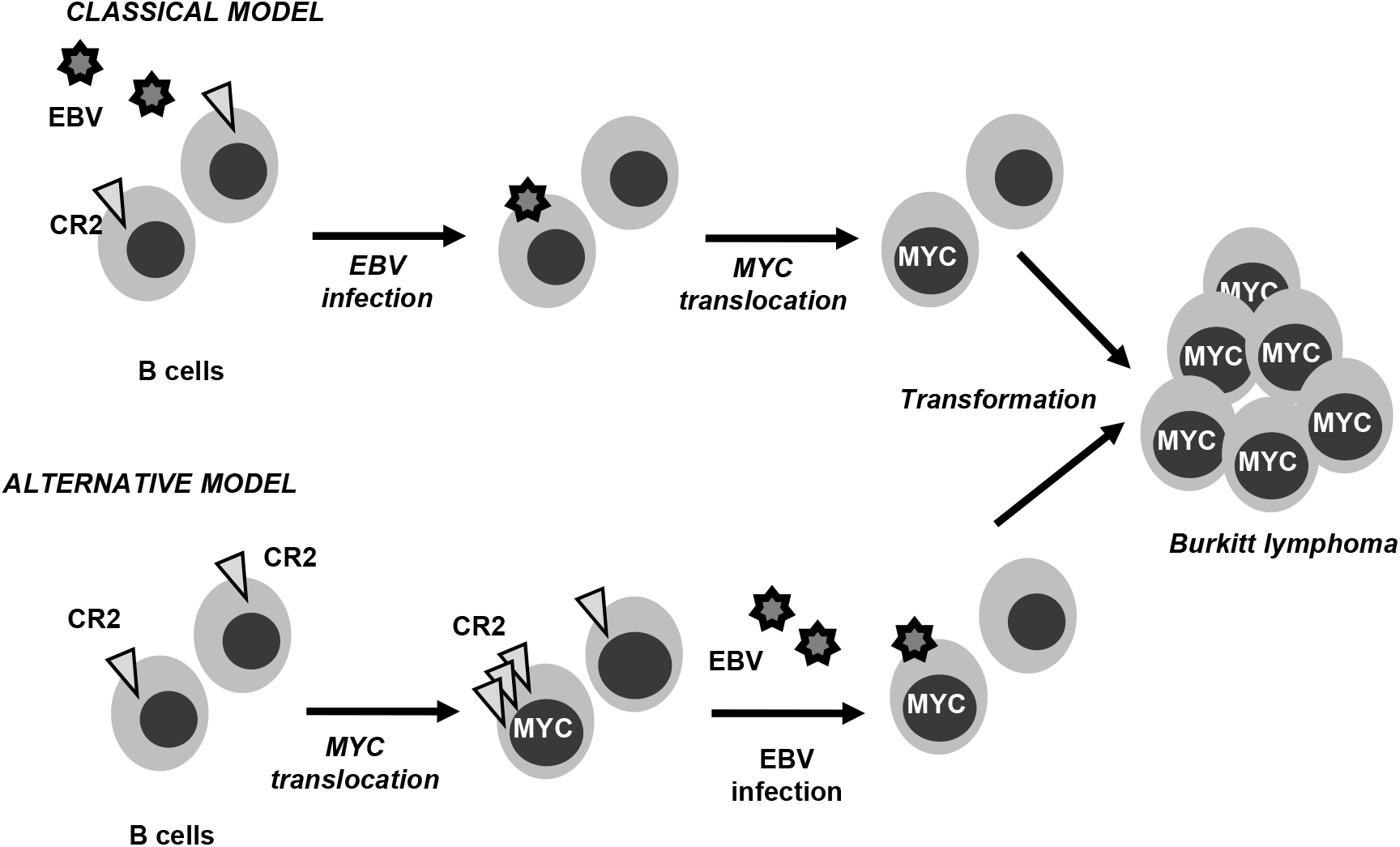
Two non-mutually exclusive models for pathogenesis of Burkitt lymphoma. In the “classical “model, EBV infection occurs first and promotes MYC translocation. The results presented here allow the proposal of a second model by which MYC translocation in tumor initiating cells leads to overexpression CR2, thus facilitating EBV infection

In view of our results it is tempting to speculate that, at least in a number of endemic BL, MYC dysregulation occurs first, facilitating subsequent EVB infection. It must be noted that our results do not contradict the canonical hypothesis (“virus first”) for BL development. Rather, they are consistent with an alternative (but not mutually exclusive) mechanism by which *CR2* upregulation by MYC would be at least partially responsible for the correlation between endemic BL and EBV infection. Both classical and alternative hypotheses are depicted in Figure 8. In accordance with the “MYC first” hypothesis, those B cells with high MYC levels (due to translocation) would show an increased density of CR2 receptors in the membrane, thus augmenting the probability of EBV infection. Consistent with this hypothesis we have shown that MYC knockdown results in decreased EBV infection whereas MYC overexpression results in increased infection in BL-derived cells. The “MYC first” mechanism is compatible with the epidemiological data and the viral clonality data, as EBV infection will co-adjuvate to lymphoma development through demonstrated mechanisms as inhibition of apoptosis and evasion of the immune surveillance, thus favoring the selection of MYC-expressing and EBV-infected cell clones. In conclusion, our results showing that CR2 is a direct MYC target gene provide new insights in the understanding of BL. Further research should be undertaken to investigate the potential of CR2 regulation by MYC as a new therapeutic approach for BL treatment.

## MATERIALS AND METHODS

### Cell culture and cell proliferation

Non-adherent cells (Raji, Jurkat, Ramos, K562, KMYCER, Kp27MYCER and B95-8) and adherent cell lines (HeLa, HEK293T) were grown in RPMI-1640 or DMEM basal media (Lonza) respectively supplemented with 10% (v/v) fetal bovine serum (Lonza), 150 μg/mL gentamycin (Lab. Normon) and 2 μg/mL ciprofloxacin. In case of KMYCER and Kp27MYCER media were also supplemented with 0.5 μg/mL puromycin and 0.5 mg/mL of G418. Viable cells were counted in hemocytometer with Trypan Blue or in a Guava cell counter (Millipore) with Guava Viacount reagent.

### Cell cycle and apoptosis analysis

Cell cycle was estimated by flow cytometry using Hoechst dye. Briefly, 10^6^ cells treated with JQ1 were harvested for each condition, centrifuged 800 x g 5 min and resuspended in previously filtered PBS (3 mM EDTA) in flow cytometry tubes. The supernatant was clarified and resuspended in PBS (3 mM EDTA) with 5 μg/mL Hoechst (33342 Thermo Fisher Scientific) and incubate 30 min at 37 °C. Apoptosis was assayed with the Annexin V-PE apoptosis detection kit (Immunostep). 10^6^ cells were harvested and washed twice with PBS (filtered with 3 mM EDTA). Cells were resuspended in Annexin V-binding buffer (10 mM Hepes/NaOH (pH 7.4). 2 μL of Annexin V-PE and 7-AAD for 100 μL were added to resuspend cells at concentration 10^6^ cells/mL. After 15 min of incubation period (in the dark) cells were resuspended in 250 μL of Annexin V-binding buffer and analyze by flow cytometry.

### RNA and DNA extraction and quantification

RNA extraction was carried out using Tri Reagent (Sigma-Aldrich). Reverse transcription (RT) was performed of total RNA extracts. 1 μg of RNA was used for reverse transcription to create complementary DNA (cDNA) with the iScript cDNA Synthesis Kit (Bio-Rad) according to manufacturer’s instructions in a total volume of 20 μl. Lately, cDNA was amplified by quantitative PCR using specific primers for the gene of interest. DNA was extracted with Qiagen DNeasy Blood & Tissue kit, following manufacturer instructions. The primers are shown in Supplementary Table 1. A PCR mix was prepared with 2xSYBR Select Master mix (Applied Biosystem), 300 nM forward and reverse primer mix and up to 35 μL of DNAse-free water (used for two 15 μL duplicate reactions) and 2 μL of each sample. PCRs were performed in the CFX Connect Real-Time PCR Detection System (Bio-Rad). Threshold cycles (Ct) were determined by default at the beginning of DNA amplification in the exponential phase. The mRNA expression of genes of interest was normalized to mRNA expression of the housekeeping gene RPS14, using the comparative Delta Ct (ΔCt) method. ΔCt=2 (Ct normalizing gene – Ct gene of interest). Reactions with water instead of cDNA were used as negative controls to detect possible amplification signals from contaminant DNA or primer dimers.

### Protein levels analysis by western blot

Roughly, 0.5-2 × 10^6^ cells were pelleted and resuspended in 100 μl of NP40 lysis buffer [50 mM Tris-HCl pH 8, 150 mM NaCl, 1 mM EDTA, 10 mM NaF, 1% NP40 (v/v), 0.1% SDS; protease (Calbiochem) and phosphatase (Sigma-Aldrich) inhibitors cocktails added immediately before use]. All the steps were performed at 4°C. Protein samples were sonicated (10 cycles of 30 s in a Bioruptor Plus Sonicator device) and finally clarified by centrifugation at 14000 rpm for 20 min at 4°C. The supernatant was transferred to a new tube and kept frozen until used. Protein quantification was carried out using Bradford Quick Start™ 1x Dye Reagent. Samples were resolved by SDS-PAGE and transferred to a nitrocellulose membrane. Primary antibodies were used diluted in TBS-T at final concentration referred in Supplementary Table 2. IRDye800/tolRDye680 secondary antibodies were used and signals were recorded with an Odyssey Infrared Imaging Scanner (LiCor Biosciences).

### Transfection and luciferase assays

Luciferase vectors (Supplementary Table 3) were transfected into hematopoietic cell lines (Raji and K562) using Amaxa Nucleofector Technology as described below. 10^6^ cells were resuspended in 100 μl Mirus Ingenio Electroporation solution and mixed with final amount 2 μg DNA plasmids (Supplementary Table 3) for each condition. The mixtures were electroporated using Amaxa nucleofector programs. Alternatively, transfection of HEK293T cells was performed using poly(ethyleneimine) (PEI) (Polysciences, Inc). Briefly, PEI and DNA were mixed in serum-free DMEM in a ratio 2.5:1 PEI:DNA (μg). Dual-Luciferase Reporter Assay System (Promega) protocol was used following manufactures’s instructions. As controls, the empty reporter pGL3 (Promega) and the *Renilla* luciferase reporter (pRLnull, Promega) were transfected. Luminescence were measured in GLOMAX Multidetection system (Promega). Firefly luminescence values were normalized against *Renilla* luminescence values used as control of transfection for each sample. Measurements were done in parallel duplicates and values were averaged. Relative light units (R.L.U.) were values related to empty vector (control) values.

### Retroviral constructs

The retroviral expression vectors were constructed by isothermal assembly of linear DNA fragments [66]. The activation-induced cytidine deaminase (AID) sequences were removed from the pMXPIE-AID retroviral vector [67] and the resulting backbone was linked to the orange fluorescent protein gene mOrange2 (from Addgene, plasmid # 45179) [68], generating the empty plasmid pMX-IRES-mOrange2. A MYC T58A gene (synthesized by IDT, Coralville, USA, using the human MYC T58A sequence as template from Addgene plasmid # 18773) [69], was linked to pMX-IRES-mOrange2 upstream to the IRES-mOrange2 cassette, generating the plasmid pMX-cMyc-IRES-mOrange2 or the empty vector without MYC, pMX-Cre-IRES-mOrange2. The lentiviral MYC expression vector (Lv224-MYC) contained the MYC-IRES-Cherry-IRES-puromycin resistance ORFs. This construct and the corresponding empty vector (Lv224) were purchased from GeneCopoeia.

### Lentivirus and retrovirus production and infection

HEK293T cells were transfected with PEI with the lentiviral packaging plasmids and the construct of interest mixed in a ratio of 1:3:4 (μg of VSV-G:psPAX2:construct) (Supplementary Table 3). Lentiviral particles-containing supernatants were collected 48 and 72 hours after transfection and stored at 4ºC. Supernatants were clarified at 1500 rpm for 10 minutes and filtered. The supernatants were mixed with PEG8000 (final concentration of 15% w:v) and incubate at 4°C for at least 6 hours. Mixture was centrifuged at 1,500xg for 30 minutes at 4°C, and the lentivirus containing pellet was resuspended in 1/200 volume of the original supernatant volume in serum-free media and stored at -80°C. For lentivirus tittering, HeLa cells were infected with increasing amounts of concentrated lentiviral particles. For Raji and Ramos infection, cells growing in RPMI with 10% FBS were plated at densities of 1.2-1.6 × 106 cells/ml and infected with concentrated lentivirus (10 – 50 μL per million cells for a MOI=3) in medium containing 5 μg/mL polybrene. The cells were incubated for 12 h and diluted 1:2 in culture medium. 36h post-infection the cells were transferred to fresh medium with puromycin (1 μg/mL), to eliminate most of the uninfected cells. After 36-48 h of incubation with puromycin, the cells were harvested for RNA and protein analysis. The empty vector pLKO and the shMYC lentiviral constructs were from Sigma Mission: TRCN0000039640, TRCN0000039642. *Lentivirus packaging vectors pCMV-*VSV-G (plasmid #8454) and psPAX2 (plasmid #12260) were from Addgene. Retroviral supernatants were produced by transient calcium phosphate transfection of NIH-293T cells with pCL-ECO (Imgenex) and pMX-Orange or pMX-MYC-Orange retroviral constructs.

### Flow cytometry and retroviral infection of mouse primary B cells

Single cell suspensions from spleens were prepared and incubated in NH4Cl buffer to lyse erythrocytes. Cells were cultured (10^6^ cells/well) in RPMI 1640 medium with 15% FBS, 2-mercaptoethanol, penicillin/streptomycin. Mature B lymphocytes were activated with LPS (20 μg/mL; Sigma-Aldrich) and IL-4 (20 ng/mL; R&D Systems) for 48 hours. Single-cell suspensions were antibody-stained in PBS with 2% FBS. Antibodies used were from BD Pharmingen (anti-CD21, anti-CD23 anti-CD69) and Biolegend (anti-B220). Spleen B lymphocytes were identified as follicular B cells (GFP^+^B220^+^CD23^+^CD21^int^), marginal zone B cells (GFP^+^B220^+^CD23^-^CD21^hi^). Mouse primary B cells were purified from spleens of 4 months old C57/BL6 wild type mice by anti-CD43 immunomagnetic depletion (Miltenyi Biotech) and cultured for 24 h in RPMI containing 10% FBS, 25 μg/mL LPS (Sigma), 10 ng/mL IL-4 (Peprotech), 10 mM HEPES (Gibco) and 50 μM 2-mercaptoethanol (Gibco) and transduced with control pMX-Orange or pMX-MYC-Orange retrovirus. CD21 expression was analyzed by flow cytometry in Orange-positive transduced cells 48 h after transduction. The antibodies are described in the Supplementary Table 2.

### Chromatin immunoprecipitation

Roughly, 2-3 × 10^7^ cells were pelleted in a 50 mL falcon, washed with PBS and centrifuged 1500 rpm 5 min. Cells were fixed with 10 mL PBS-1% formaldehyde, rotating for 10 min and the fixation was blocked with 125 mM Glycine for 5 min keeping the cells in rotation. Supernatant was clarified at 1500 rpm for 5 min and washed twice with PBS, before storage at -80 °C. Cells were lysed with 1.2 mL of lysis buffer (20 mM Tris HCl pH 8.0; 2 mM EDTA; 0.7% SDS; 1% Triton X-100; 150 mM NaCl; H_2_O), adding protease and phosphatase inhibitor immediately before used and led the cells 10 min on ice before sonicate on Bioruptor for 10-13 cycles (30 s ON, 30 s OFF). Supernatants were clarified at 14000 rpm at 4°C and collected in a new 1.5 mL eppendorf and stored at -80 °C. The lysate were diluted 7 times in Dilution buffer (protease inhibitor cocktail 1:100, added immediately before use). We also added also 2-3 μg of our antibody to the lysate and incubate the mix rotating at 4°C overnight. After antibody incubation, 30 μl of Dynabeads Protein G (Invitrogen) were added and incubated them for 30 minutes to 2 hours, rotating at 4°C. Previously, the Dynabeads were blocked for 30 min with Sperm DNA salmon (Invitrogen) (1 μg/ mL for each sample). After incubation with the chromatin samples, the beads were washed in cycles of 5 minutes with Low Salt Buffer (0.1% SDS; 1%Triton X-100; 2 mM EDTA; 20 mM Tris-HCl pH 8; 150 mM NaCl), High Salt Buffer (0.1% SDS; 1%Triton X-100; 2 mM EDTA;20 mM Tris-HCl pH 8; 500 mM NaCl), LiCl Buffer (0.25 M LiCl; 1%NP 40 (IGEPAL); 1% Sodium Deoxycholate; 1 mM EDTA; 10 mM Tris-HCl pH 8), two washes with TE Buffer (10 mM Tris HCl pH 7.5;1 mM EDTA) and resuspended in 200 μL of Elution Buffer (1%SDS; 50 mM Tris-HCl pH 7.5). Samples were incubated with Elution buffer shaking at 65°C for 30 to 40 minutes, then they were des-crosslinked with 6 μL RNAsa (10 mg/mL) and 12 μL NaCL 5M and incubated shaking at 65°C, overnight. Once separated chromatin from proteins we added 2 μM EDTA 0.5 M, 4 μL Tris pH 6.5 1 M and 1 μL Proteinase K (20 mg/ mL, Thermo Fisher Scientific) and incubated shaking for 3 hours at 45°C. DNA were purified with Qiaquick PCR purification kit (Qiagen), as it has been described above. We performed qPCR with 4 μL of each sample. Primers were designed with Primer3 software as mentioned above and primer sequences are detailed in Supplementary Table 1.

### EBV infection

B95-8 cells were seeded to confluence 0.5-0.9 × 10^6^ cells per mL in a flask with 100 mL of RPMI 10% FBS. Let the cells growing for 3 weeks without changing or adding media. After 3 weeks the supernatants were collected, filtered with 0.45 μm filter and storage in 5 mL tubes. Ramos cells were infected, with lentiviral particles, carrying shRNA against *MYC*, for 48 h and selected with puromycin for further 48 h. The remaining cells were collected and plated in a T12 and 500 μL of these cells were seeded with 500 μL of the EBV virus supernatant. Cells were harvested at different time points and pelleted for RNA and DNA extraction. RNA and DNA were extracted with Qiagen kits following manufacturer instructions. 50 ng of DNA were quantified by qPCR analysis. *CR2* and *LDHA* promoter primers were used as loading control and primers that recognize DNA viral latent genes (Supplementary Table 1).

## Acknowledgements and Funding

The work was supported by grants PID2020-115903GB-100 to J.L. and M.D.D., RTI2018-095673-B-I00 to J.R.R., PID2019-107551RB-I00 to V.G-Y., and PID2019-106773RB-I00 to A.R.R, funded by MCIN/AEI/10.13039/501100011033/, Spanish Government, and by “FEDER, Una manera de hacer Europa”, European Union, and by La Caixa HR17-0244 grant to A.R. J.L-P. and L.Q. were recipients of F.P.U. program and Universidad de Cantabria fellowships, respectively. We are grateful to M. Dolores Delgado for insightful comments and critical reading of the manuscript, and Maria Aramburu and Patricia Arribas for technical help.

