## Supplemental tables and figures for "MYC directly transactivates CR2/CD21, the receptor of the Epstein-Barr virus, and enhances the viral infection of Burkitt lymphoma cells"

**Ester Molina et al.**

#### SUPPLEMENTARY TABLES AND FIGURES

**Supplementary Table S1.** Primers used in PCR**.** All correspond to human genes**.** The primers are written in the 5'→3' direction

| Gene | Forward | Reverse | Product Size (bp) | Use |
|---|---|---|---|---|
| CR2 | CCGACACGACTACCAACCTG | GACAATCCTGGAGCAATGGA | 115 | RT-qPCR |
| CR2 +325 bp | ATAAACCGCCTGGTCCTGAT | CCTCATGCAGTGGGTAGGTT | 235 | ChIP |
| CR2 +481 bp | AGGATTCTGCAGGTGCTCAT | TGCTTGGGAACCAGGTCTTA | 196 | ChIP |
| CR2 +141 bp | GGGTTTTCTTGGCTCTCGTC | ATTGCAGTGGTCCCTCAAAG | 188 | ChIP |
| CR2 -35 bp | GTGTGCGCTCAGAACTAGCA | CGACGAGAGCCAAGAAAACC | 197 | ChIP |
| CR2 -198 bp | AGTGTAGTGGGTTGCGTGGT | GCGGGCCCTTAAATAGTGTC | 212 | ChIP |
| CR2 -525 bp | CATGCAGAGAATCTGGGTGA | GTGCCATGCTGTGTTATGCT | 246 | ChIP |
| CR2 +2.1 Kb | GCATTCCTTGGAGAAACAGC | CAGTGGGAGCCCTCAAAATA | 153 | ChIP |
| CR2 +2.2 Kb | GGGTGCGGAAACAATGATAC | CAGTGGGAGCCCTCAAAATA | 243 | ChIP |
| CR2 Exon9 | CAAGAAAGAGGCACCTGGAG | CAGCTTTGCAACGAATCTGA | 199 | ChIP |
| CR2 promoter | GCTTGTCCCACCCTCACC | GTTCCATCTTCCAGCGGATA | 221 | ChIP |
| EBNA1 | GGTTCCAGCCAGAAATTTGA | CTCGCCTGAGGTTGTAAAGG | 151 | PCR |
| LDHA | TCCTGACTCAGGCTCATGGC | AGACAACCGACCGGCAGA | 103 | RT-qPCR |
| LMP1 | CGCCTAGGTTTTGAGAGCAG | GATGAACACCACCACGATGA | 154 | PCR |
| MYC | TCGGATTCTCTGCTCTCCTC | CCTGCCTCTTTTCCACAGAA | 157 | RT-qPCR |
| CDKN1B/p27 | CCGGCTAACTCTGAGGACAC | AGAAGAATCGTCGGTTGCAG | 120 | RT-qPCR |
| RPS-14 | TATCACCGCCCTACACATCA | GGGGTGACATCCTCAATCC | 135 | RT-qPCR |

**Supplementary Table 2.** Antibodies used in this work. IB, immunoblot; FC, flow cytometry.

| **Antigen** | **Reference and specie** | **Concentration, Use** |
|---|---|---|
| MYC | N262, Sc-764, rabbit polyclonal Santa Cruz | 1:1000 IB<br>1:40, FC |
| MYC | 9402, , rabbit polyclonal Cell Signaling | 1:3000, IB |
| MYC | D84C12, mouse monoclonal Cell Signaling | 1:200, FC |
| CR2/CD21 | C-20 Santa Cruz Biotech. sc-7025, goat polyclonal, | 1:1000, IB |
| CR2/CD21 | anti-CD21-APC ref 558658, Biolegend | 1/100, FC |
| CD23 | anti-CD23-PE ref 553139, Biolgened | 1/100, FC |
| CD69 | anti-CD69-bio-APC ref 553235) Biolegend | 1/100, FC |
| B220 | anti-B220-PE-Cy7, ref 103222 Biolegend | 1:200 FC, |
| β-Actin | I-19, sc-1616, goat polyclonal, Santa Cruz Biotech. | 1:1000, IB |
| β-Actin (C-4) | Sc-47778, mouse, Santa Cruz Biotech. | 1:3000, IB |
| Cyclin A2 (H-432) | Sc-239, rabbit polyclonal, Santa Cruz Biotech. | 1:1000, IB |
| PARP-1 (H-250) | Sc-7150, rabbit polyclonal, Santa Cruz Biotech. | 1:1000, IB |
| CR2/CD21 FITC | Rat SD IgG2b,K, clon 7G6, BD Pharmingen | 1:200, FC |
| CD23 biotin | Rat (LOU/M) IgG2a,K, clon B3B4, BD Pharmingen | 1:200, FC |
| PE Streptavidin | BD Pharmingen | 1:400, FC |
| CD19 APC eFluor780 | Rat IgG2a,K, clon eBio1D3 (1D3), BD Pharmingen | 1:100, FC |

**Supplementary Table 3.** Plasmids used in this work

| **Plasmids** | **Construct** | **Origin** |
|---|---|---|
| pmaxGFP | GFP gene | Amaxa |
| pGL3 | *Firefly* sp. luciferase reporter gene with no promoter region | Promega |
| CR2-Luc | -315bp CR2 promoter in pGL3 | [52] |
| Ebox1Mut-Luc | -315bp CR2 promoter with Ebox1 deletion in pGL3 | [52] |
| Ebox2Mut-Luc | -315bp CR2 promoter with Ebox2 deletion in pGL3 | [52] |
| 4Ebox-Luc | *Firefly* sp. Luciferase reporter gene regulated by 4 E-box sequences | [70] |
| pRL-null | *Renilla* sp. luciferase reporter gene regulated by the T7 promoter | Promega |
| pCMV-VSV-G | VSV-G gene encoding enveloped lentiviral protein | Addgene, plasmid #8454 |
| psPAX2 | GAG and POL genes enconding packaging lentiviral proteins | Addgene, plasmid #12260 |
| pLKO.1 control | Empty vector | Sigma- MISSION |
| pLKO.1 shMYC | Short hairpin RNA TRCN0000039640 against human MYC | Sigma- MISSION |
| pLKO.1 shMYC | Short hairpin RNA TRCN0000039642 against human MYC mRNA | Sigma- MISSION |
| pLKO.1 shCR2 | Short hairpin RNA TRCN0000057113 against human CR2 | Sigma- MISSION |
| pLKO.1 shCR2 | Short hairpin RNA TRCN0000057114 against human CR2 | Sigma- MISSION |
| pMX-cMyc-IRES-mOrange2 | Retroviral vector carrying human MYC T58A ORF and mOrange2 ORF separated by an IRES sequence | This work |
| pMX-IRES-mOrange2 | Retroviral vector carrying the mOrange2 gene and an IRES sequence | This work |
| Lv224-MYC | Lentiviral vector expressing MYC-IRES-Cherry-IRES-puromycin resistance gene | Genecopoiea |
| Lv224 | Lentiviral vector expressing Cherry-IRES-puromycin resistance gene | Genecopoiea |

### Supplementary Figure S1

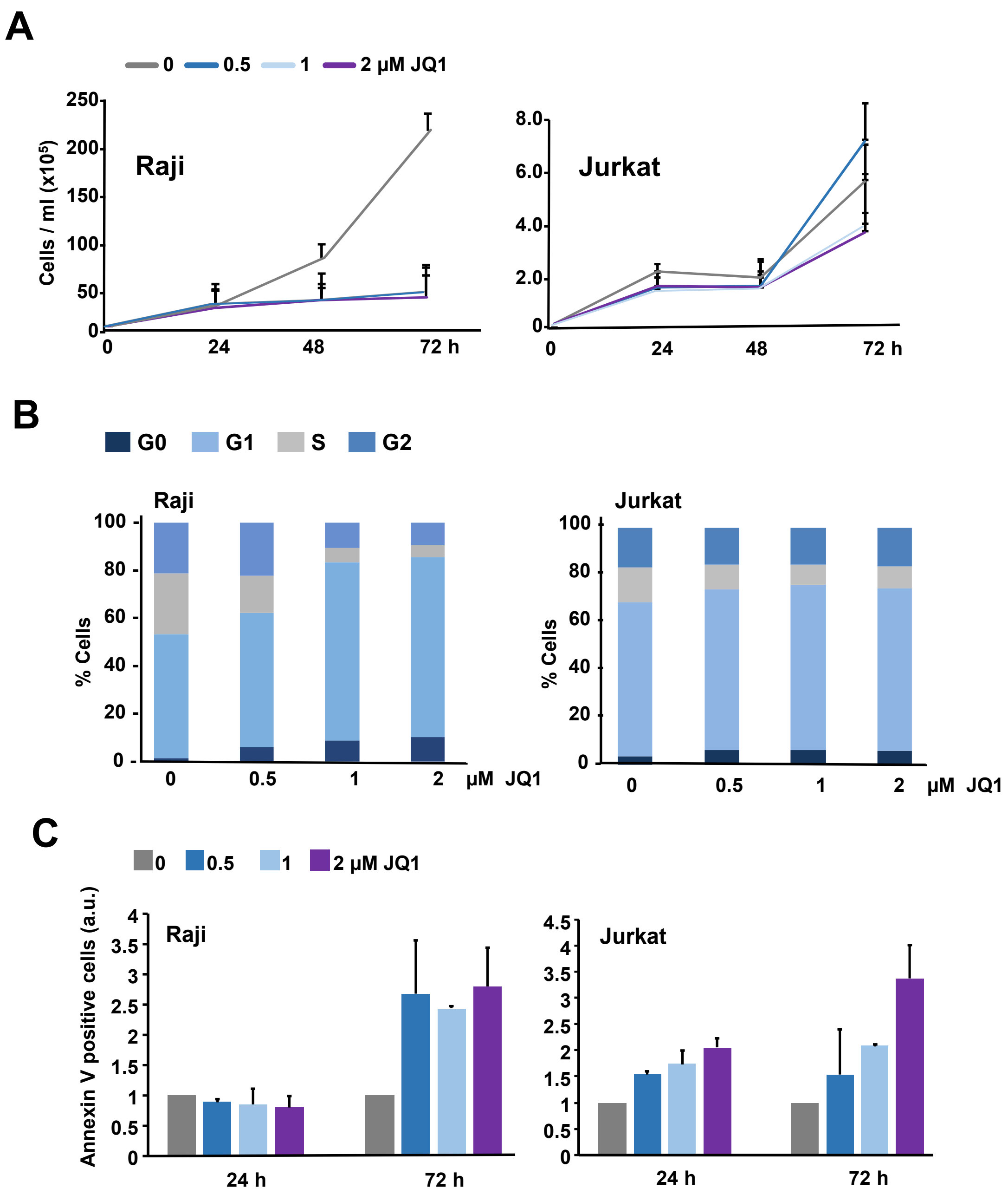


**Supplementary Figure 1. Effect of JQ1 on proliferation and apoptosis of Raji and Jurkat cells. (A)** Proliferation of Raji and Jurkat cells treated with JQ1 at the indicated concentrations for up to 72 h, estimated by cell counting. Data represent mean values ±S.D. (n = 3). (**B**) Cell cycle distribution of Ramos and Jurkat cells treated for 48 h with JQ1 at the indicated concentrations. Cells were stained wit iodide propidium and analyzed by flow cytometry. Data are mean values from two independent experiments ±SD. (**C**) Apoptosis of Raji and Jurkat cells assayed by annexin V binding and measured by flow cytometry. Data represent means values ±S.D. (n=2). Graphs represent the fraction of positive cells normalized to the value of untreated cells.

### Supplementary Figure S2

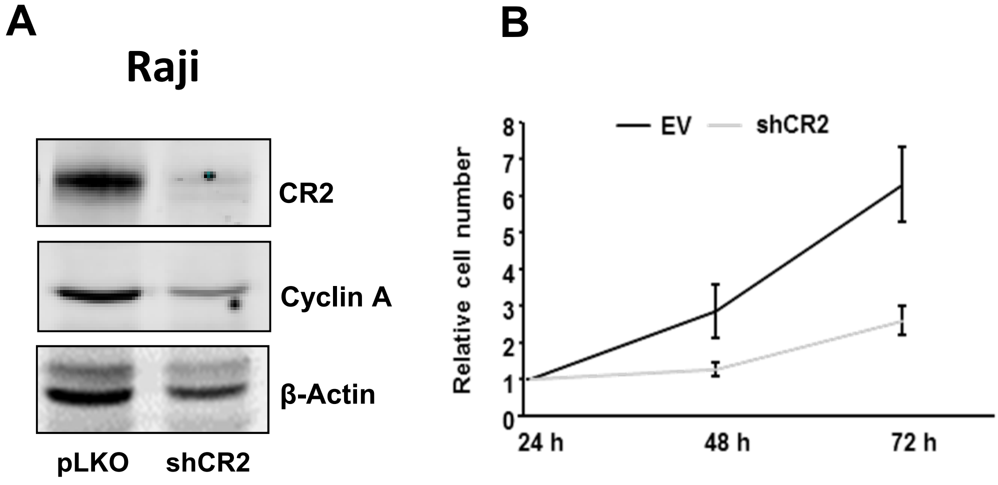


**Supplementary Figure S2.** Silencing of CR2 impairs Raji cell proliferation. (**A**) Cells were infected with a mixture of two lentiviral vectors encoding for CR2 short-hairpin constructs as well as the pLKO empty vector (EV). After 36 h puromycin was added (1 µM final concentration) and the cells were further incubated for 72 h. Cells were then harvested and the indicated proteins analyzed by immunoblot. Expression of the indicated proteins 48 h after the addition of puromycin. (**B**) Viable cell densities, as determined by trypan blue at the indicated time points

### Supplementary Figure S3

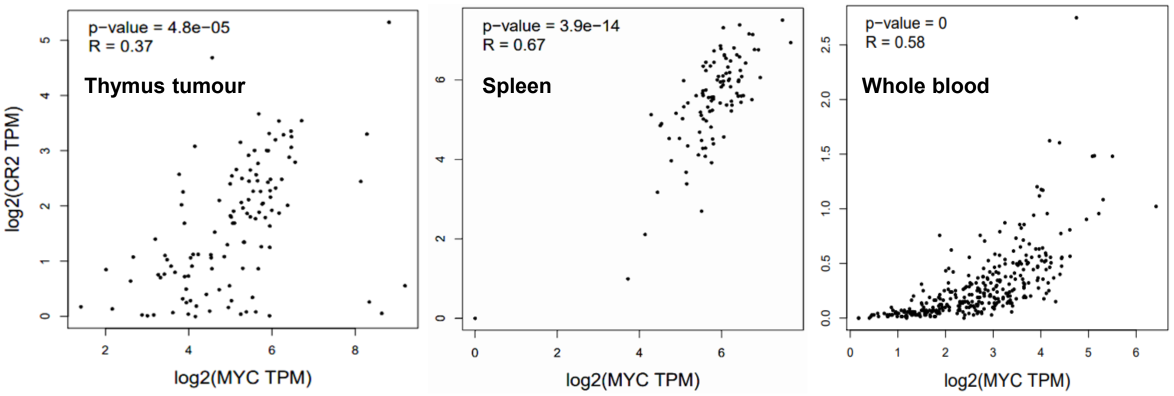


**Supplementary Figure S3.** Expression correlation between mRNA levels of MYC and CR2 in human thymus tumours (TCGA database), normal spleen (TCGA database) and spleen (GTEx database) according to the Gene Expression Profiling Interactive Analysis (http://gepia.cancer-pku.cn/) using the GTEx tissues database. R, Pearson correlation coefficient

### Supplementary Figure S4

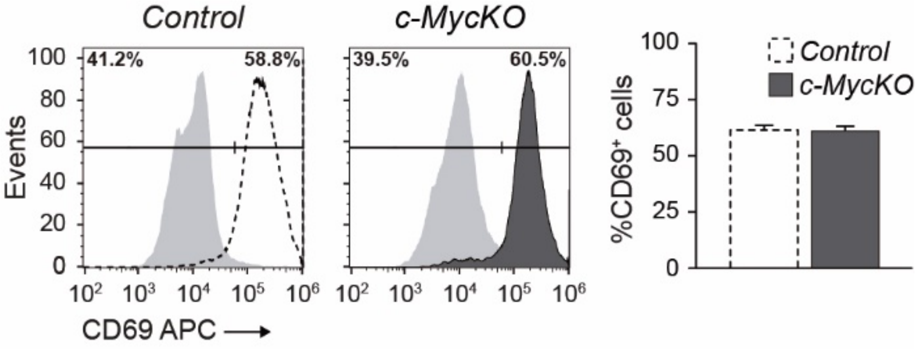


**Supplementary Figure S4**. CD69 staining of mouse splenic B cells. Mature B lymphocytes from the spleens of *Myc*$^{flox/flox}$*;Max*$^{flox/+}$*;cd19*$^{cre/+}$*;Rosa26*$^{gfp/gfp}$ (MycKO, *n*=3) homozygous and *Myc*$^{flox/+}$*;Max*$^{flox/+}$*;Cd19*$^{cre/+}$*;Rosa26*$^{gfp/gfp}$ heterozygous control mice (*n*=3) were activated with LPS and IL-4 for 48h. GFP$^{+}$ cells (*Myc*-depleted B lymphocytes) were gated and surface expression of CD69 was analysed by flow cytometry. Absolute numbers of CD69$^{hi}$ are shown, ±S.D. (n=3). ***$p<0.001$.

### Supplementary Figure S5

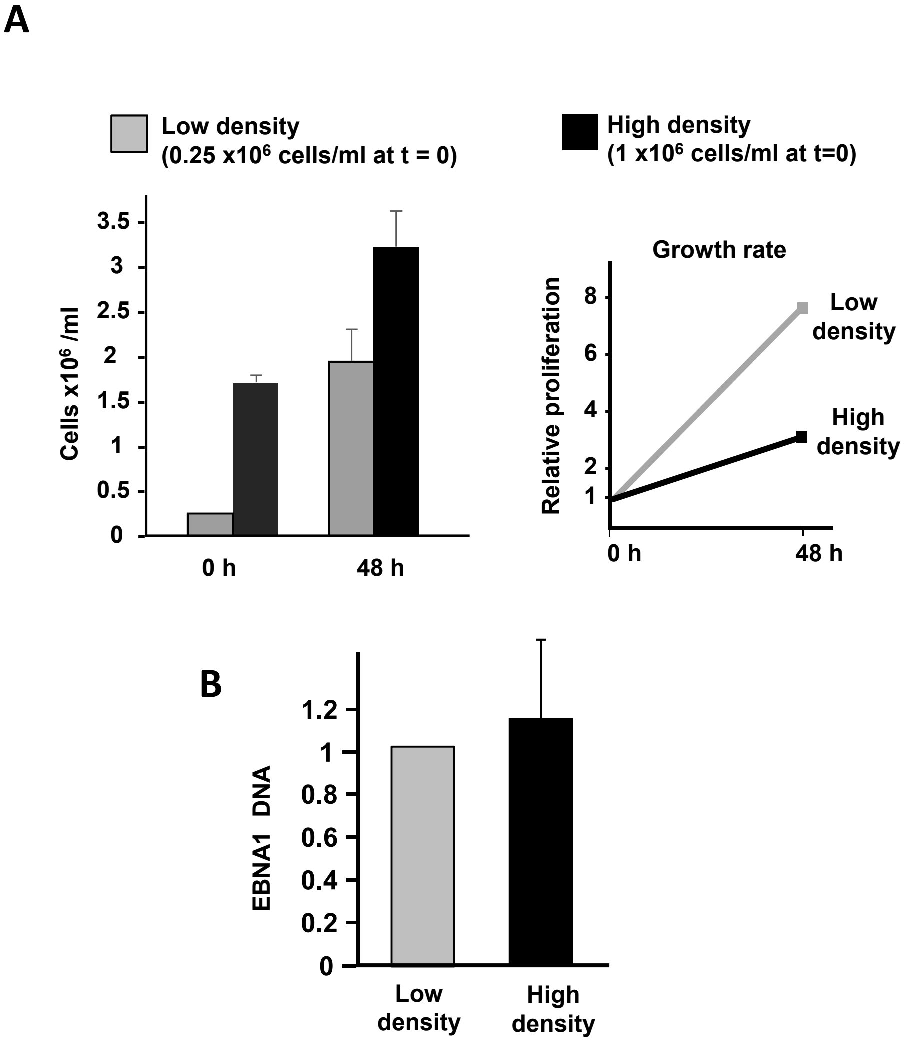


**Supplementary Figure S5. Ramos cells proliferation rate does not affect the infection by EBV.** (**A**) Cell proliferation rate of the cells infected at high and low density cultures. Ramos cells were grown until a density of 0.5 x $10^5$ cells/ml /low density) or 2 x $10^6$ cells/ml (high density), infected with EBV (1:1 vol de B95-8 supernatant) and further incubated for 48 h. The cells were then harvested, washed and the total DNA extracted, The cell densities are shown at the left graph and the relative growth rate at the right graph (n = 5). (**B**) The levels of viral genomic DNA were determined by PCR with primers of the EBNA1 gene. Data was normalized to the mean DNA levels of LDH and CR2 genomic DNA (n=5)

### Supplementary Figure S6

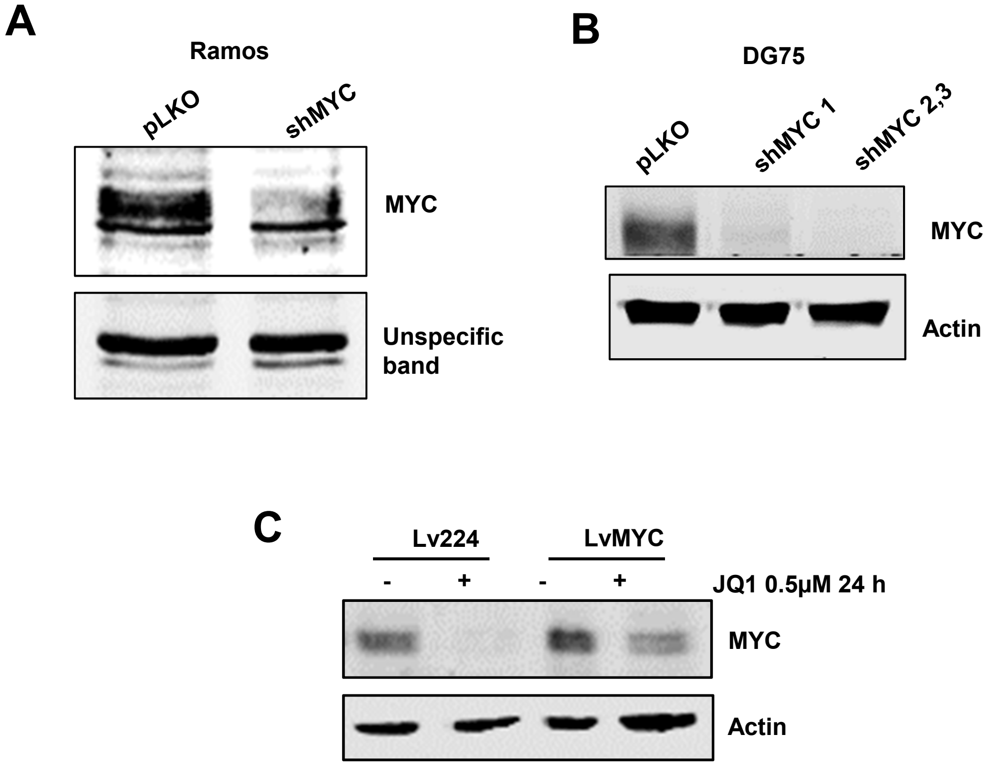


**Supplementary Figure S6**. Silencing of MYC in BL cells. (**A**) Ramos cells where infected with a mixture of shMYC1 and shMYC2 lentivirus as described in Methods. The cells were harvested, lysed and the expression of MYC protein was analyzed by immunoblot (B) DG75 cells were infected by shMYC1 and shMYC2 lentivirus as described in Methods and the expression of MYC protein was analyzed by immunoblot. The expression of β-actin was analyzed as loading control. (**C**) DG75 cells were infected with lentivirus expressing MYC (Lv224-MYC) and the empty vector, selected for 10 days with puromycin 0.75 µg/ml and treated with JQ1 (0.5 µM) for 24 h to suppress endogenous MYC expression. The expression of MYC and β-actin (as loading control) was analyzed by immunoblot.
